# Discovery and validation of the binding poses of allosteric fragment hits to PTP1b: From molecular dynamics simulations to X-ray crystallography

**DOI:** 10.1101/2022.11.14.516467

**Authors:** Jack B. Greisman, Lindsay Willmore, Christine Y. Yeh, Fabrizio Giordanetto, Sahar Shahamadtar, Hunter Nisonoff, Paul Maragakis, David E. Shaw

## Abstract

Fragment-based drug discovery has led to six approved drugs, but the small size of the chemical fragments used in such methods typically results in only weak interactions between the fragment and its target molecule, which makes it challenging to experimentally determine the three-dimensional poses fragments assume in the bound state. One computational approach that could help address this difficulty is long-timescale molecular dynamics (MD) simulation, which has been used in retrospective studies to recover experimentally known binding poses of fragments. Here, we present the results of long-timescale MD simulations that we used to prospectively discover binding poses for two series of fragments in allosteric pockets on a difficult and important pharmaceutical target, protein-tyrosine phosphatase 1b (PTP1b). Our simulations reversibly sampled the fragment association and dissociation process. One of the binding pockets found in the simulations has not to our knowledge been previously observed with a bound fragment, and the other pocket adopted a very rare conformation. We subsequently obtained high-resolution crystal structures of members of each fragment series bound to PTP1b, and the experimentally observed poses confirmed the simulation results. To the best of our knowledge, our findings provide the first demonstration that MD simulations can be used prospectively to determine fragment binding poses to previously unidentified pockets.

---

Fragment-based drug discovery (FBDD) aims to identify small molecules of low molecular weight (fragments) that bind a biomolecule of interest. Such fragments typically bind weakly themselves, but are used as starting points for the design of more potent compounds. FBDD often involves the experimental evaluation of binding for many fragments (a process known as “fragment screening”). Many pharmaceutical companies use fragment screening in their early-stage drug discovery pipelines to identify new chemical matter, complement high-throughput screening, or refine the understanding of a particular binding site.^1,2^ Such laboratory fragment screens commonly make use of biophysical assays, including nuclear magnetic resonance spectroscopy (NMR), X-ray crystallography, and surface-plasmon resonance (SPR)^1,2^ to identify fragments that bind their intended target (hits). These efforts have already resulted in six approved drugs that were designed starting from a fragment hit.^3^

To validate a fragment hit and improve its drug-like properties, it is often important to understand the binding site and pose of the fragment.^1,2^ It remains challenging, however, to do so using experimental means: By definition, fragments have low molecular weight (typically <300 Da) and associate with their target with millimolar to micromolar affinity,^1,2^ and the various biophysical assays used to identify fragment hits typically have low signal-to-noise ratios for low-molecular-weight or weak binders, meaning that these techniques often give inconclusive, and in some cases, contradictory results about which fragments interact with the target. Indeed, a requirement that two biophysical techniques agree with each other often results in the elimination of the majority of all fragment hits.^4^

Molecular dynamics (MD) simulation can model the motions of all the atoms of a biomolecular system of interest, and is thus a potentially powerful tool to identify the binding modes of fragment hits. The most direct MD approach is to simulate the pharmaceutical target of interest and the surrounding solvent containing one or more fragments, in order to observe the diffusion of the fragments in solution and their eventual association with the target. Such an approach is a potentially powerful one for identifying binding modes, because it does not require any a priori knowledge of the bound conformation, in particular the location of the binding site, information which is often unavailable in practice. MD simulations have already been used for the related problem of identifying new interactions that could be used to enhance ligand-binding affinity,^5,6^ and several MD simulation studies have been used to reproduce the previously known crystallographic binding poses of ligands to their target proteins.^7–15^ To date, however, we are not aware of any published MD simulation study that has predicted the pose of a ligand in a previously unknown binding site, and that has subsequently been verified by X-ray crystallography experiments.^16^

Here, we present the results of long-timescale, all-atom MD simulations that we used to discover the binding sites and poses of two fragments to protein-tyrosine phosphatase 1b (PTP1b), an important, but difficult pharmaceutical target. Prior to our simulations, no structures were available for either of these fragments bound to PTP1b. In our simulations, we observed repeated binding to and unbinding from their respective bound poses in two distinct allosteric sites (pockets 1 and 2). For the fragment that bound in pocket 2, we observed the rearrangement within the pocket of two phenylalanine side chains adjacent to the fragment that led to a PTP1b conformation rarely observed in crystal structures. We then obtained high-resolution crystal structures of PTP1b bound to each fragment and found that these crystal structures confirmed the poses observed in simulation. To provide additional evidence that these binding poses are not artifacts, we synthesized fragments that were chemically closely related to the hits and obtained high-resolution crystal structures of PTP1b bound to these related fragments in poses that overlapped with the poses of the original hits. These results demonstrate, for the first time, MD simulations being used in the successful prediction of the previously unknown binding sites and poses of fragments to their target proteins.

PTP1b—a negative regulator of the insulin and leptin pathways and of unfolded protein response in the endoplasmic reticulum—has been a prototypical single-domain protein for experimentation with technologies for allosteric-binding-site discovery. It was previously considered a potentially attractive target for treating type II diabetes and obesity, though concerns regarding skin lesions in animal models, and difficulties encountered in finding potent and selective compounds that are active in the cell, have limited recent efforts in PTP1b drug discovery.^17–19^ PTP1b is also involved in the development of Her2-positive breast cancers, and a product derived from dogfish liver that is currently an early-stage clinical candidate for this indication is believed to act on PTP1b by an allosteric mechanism.^20^ Because of the conserved and very polar nature of the active site of PTP1b, it is widely believed that only allosteric inhibitors of PTP1b have a realistic chance of becoming useful drugs, and for this reason PTP1b has become a workhorse for methods that attempt to elucidate allosteric pockets.^21–27^

As part of our in-house drug discovery efforts, we ran an SPR screen of two 384-well plates of the Maybridge library (Thermo Fisher Scientific) and identified two fragments (referred to here as DES-4799 and DES-4884) that showed stronger binding to PTP1b than to five off-target proteins that are structurally unrelated to PTP1b (see Supplementary Information). We then used MD simulations of these two fragments in the presence of PTP1b to predict their binding sites and poses. The fragments were not subject to any artificial biasing forces, and were free to diffuse about the simulation box and sample potential binding modes with PTP1b. (See SI for further details regarding the setup of these MD simulations.)

DES-4799, an anilinic fragment, bound reversibly in our simulations to pocket 1, a novel allosteric binding site located diametrically opposite from the active site. Fig. 1 shows that this aniline reversibly associated with a small number of sites on the surface of the protein over the course of a 100-μs MD simulation (see Movie S1). At the fragment concentration in this simulation, the fragment was unbound for the majority of the simulation time, suggesting an affinity >3 mM. The single dominant binding site was defined by residues Ser80, Ser205, His208, and Gly209; the conformation of these residues was stabilized upon binding (Fig. S3). The location of this binding site, along with those of three additional minor binding sites, are highlighted in Fig. 1b. In the main site, the aniline bound in three different orientations (Fig 1c), with a strong preference for one of the three poses (see histogram in Fig 1a).

**Figure 1.**
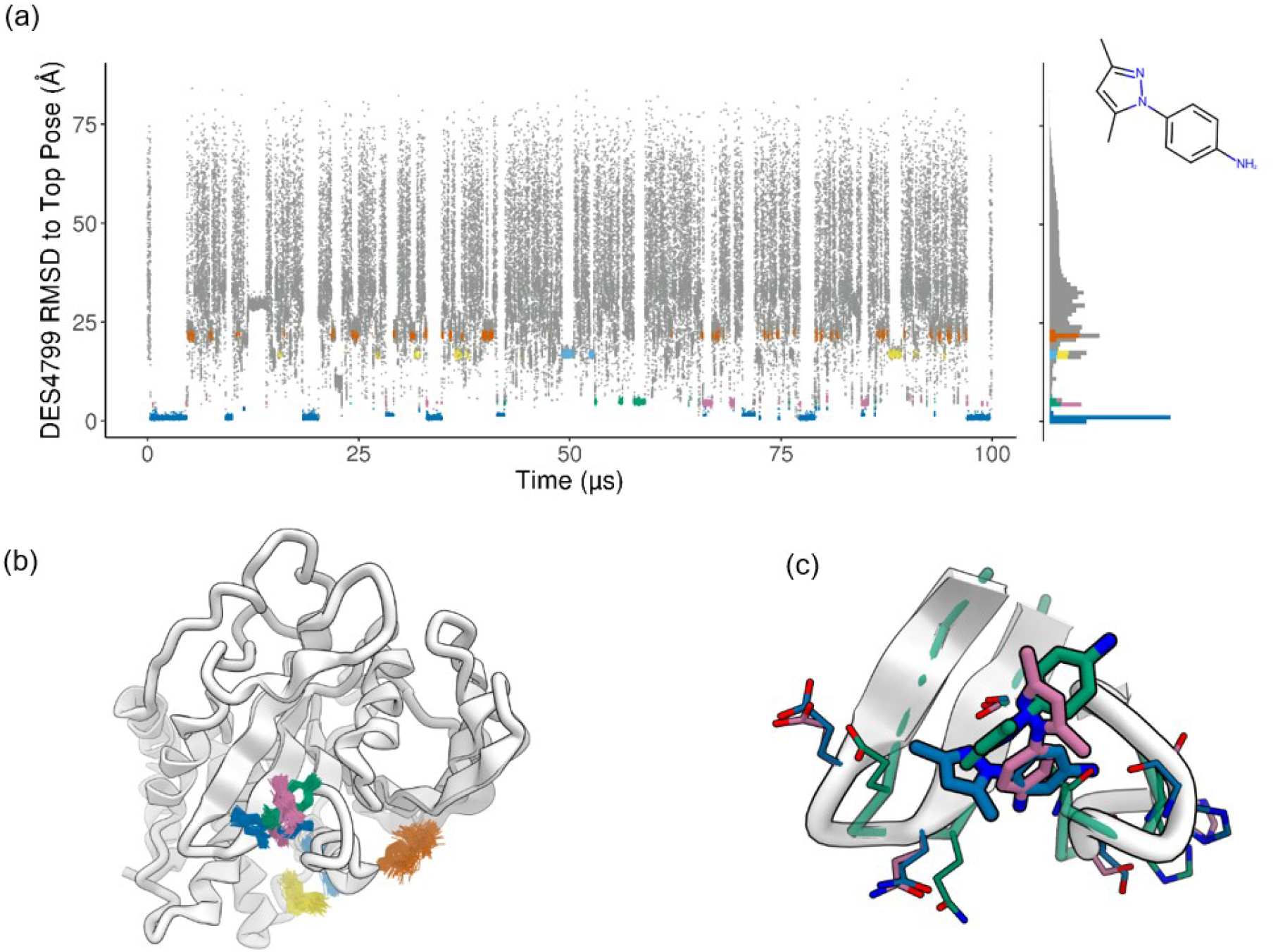
Summary of a representative 100-μs MD simulation of DES-4799. (a) The time series and probability densities of the RMSD of the small molecule to the dominant binding pose with the protein backbone aligned to the initial structure of the simulation. The data from the top six binding clusters of the fragment density are shown in color. (b) The locations of the top binding clusters on the surface of the protein, represented by 100 snapshots closest to their respective centroids. (c) The orientation of the fragment in the three clusters that bind in the main pocket.

DES-4884, a piperazinylquinoline fragment, bound in our simulations to pocket 2, which overlaps with the allosteric pocket discovered by Sunesis.^22^ In our simulations, however, binding of the fragment involved a rearrangement of aromatic side chains not previously observed in PTP1b X-ray structures. Fig. 2 shows that DES-4884 reversibly bound to its main binding site several times in two different orientations. It also visited a small number of other binding sites, but spent at least an order of magnitude more time at the primary binding site than at any of these secondary sites (see Movie S2). The simulated binding affinity to the main site was ~1 mM. Fig. 2b shows the locations of the secondary PTP1b binding sites visited by the piperazinylquinoline fragment; the site colored in orange overlaps with a pocket recently identified by Keedy et al.^23^

**Figure 2.**
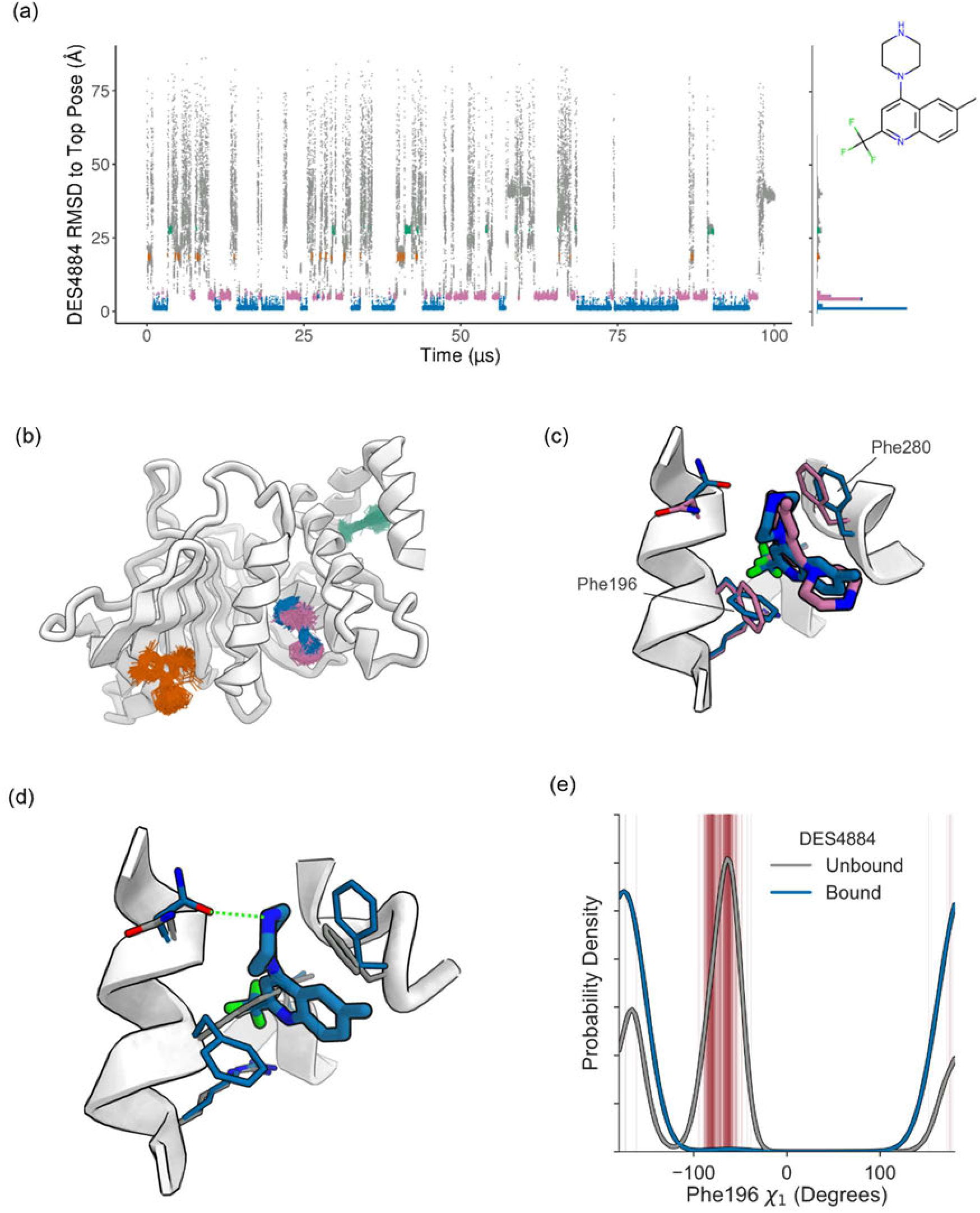
Summary of a representative 100-μs MD simulation of DES-4884. (a) The time series and probability densities of the RMSD of the small molecule to the dominant binding pose with the protein backbone aligned to the initial structure of the simulation. The data from the top four binding clusters of the fragment density are shown in color. (b) The locations of the top four binding clusters, represented by 100 snapshots closest to their respective centroids. (c) The orientation of the fragment in the two clusters that bind in the main pocket. (d) The main binding pose of DES-4884; the fragment forms a hydrogen bond with Asn193 and the side chains of Phe280 and Phe196 swing out (shown in blue) compared to their apo orientations (shown in gray). (e) The density of the χ_1_ angle of Phe196 during the snapshots in which the fragment is bound to the main pocket (blue) and the snapshots in which the fragment is not bound (gray). The red lines highlight the χ_1_ angle of Phe196 in all crystal structures of PTP1b published in the PDB, including a line for each independent chain or alternate conformation.

Fig. 2c shows the two distinct binding orientations of DES-4884. In the dominant orientation, the piperazine formed a hydrogen bond with Asn193. In both orientations, the pocket rearranged to a conformation distinct from most previously reported crystal structures: Both Phe280 (top right) and Phe196 swung out to open a pocket that allowed for hydrophobic contacts with the quinoline core (Fig. 2d). Phe196 does not adopt the swung-out configuration in the vast majority of crystal structures published to date (Fig. 2e). The only published structures in which Phe196 adopts the swung-out configuration can be explained by crystal contacts,^24^ nearby mutations,^28^ or fragment binding.^23^ In simulation, the side chain remained swung out while the fragment was bound, and then rearranged to the dominant apo configuration while the fragment was not bound (Fig 2e) (see Movie S3).

Following binding characterization by MD simulation, we validated the predicted binding poses by obtaining high-resolution crystal structures of PTP1b in complex with each of the two fragments. To further validate the binding poses we identified for these fragments, we obtained additional high-resolution crystal structures of two variants of DES-4799 and one variant of DES-4884 that recapitulated their respective binding modes. Fig. 3 shows that the dominant binding poses from simulation very closely match the high-resolution crystal structures, including structurally important details such as the location of water molecules bound to each of the fragments.

**Figure 3.**
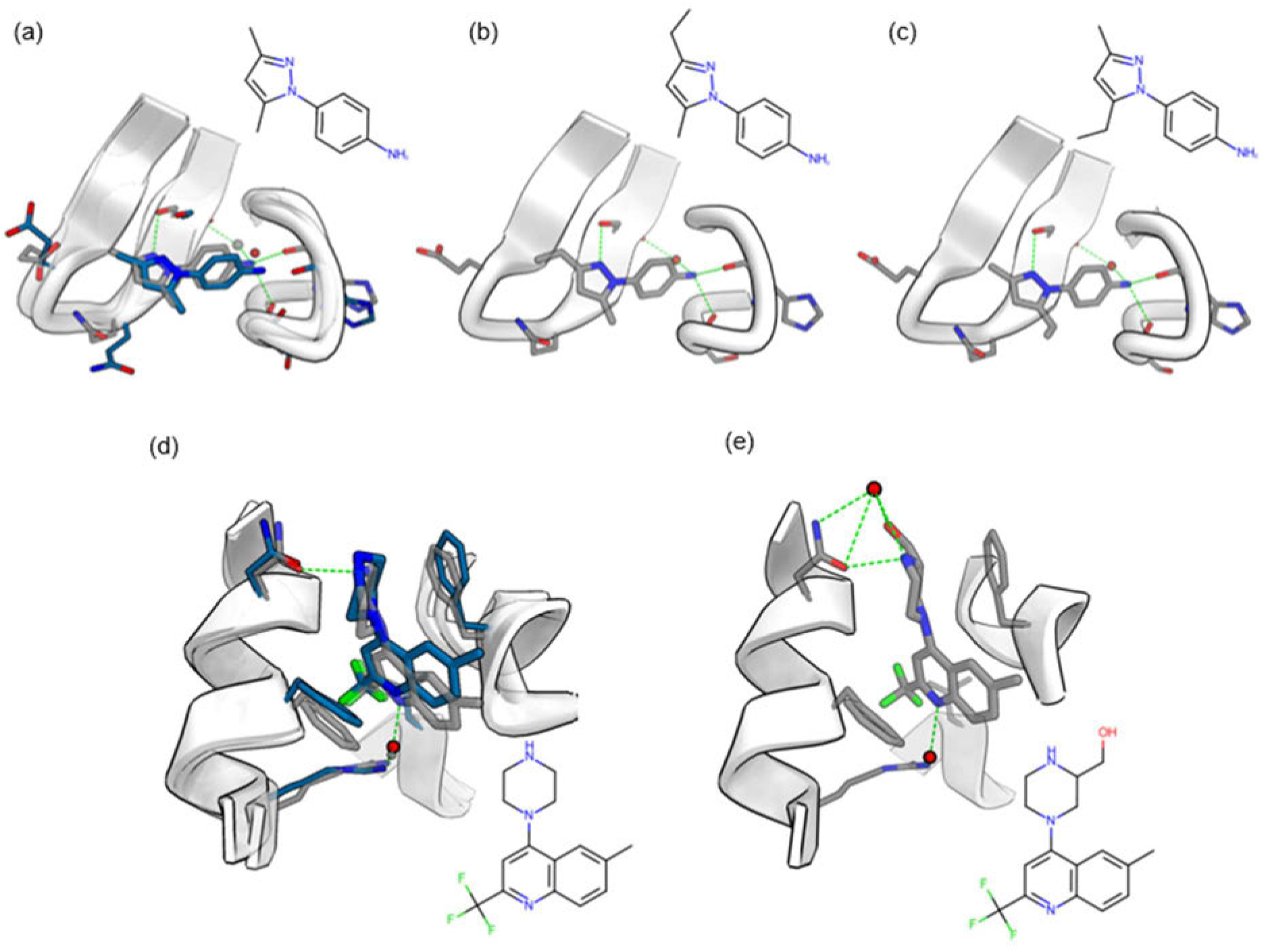
X-ray structures of DES-4799, DES-4884, and respective variants bound to PTP1b. (a) The top binding pose of DES-4799 predicted by MD simulation (blue) overlaid on its high-resolution X-ray crystal structure (gray). The oxygen atom of water is represented by a red sphere (simulation) or by a grey sphere (crystal structure). (b) The binding mode of the crystal structure of DES-5742, a single-methyl variant of DES-4799. (c) The binding mode of the crystal structure of DES-5743, a single-methyl variant of DES-4799. (d) The top binding pose of DES-4884 and pocket conformational changes predicted by MD simulation (blue) overlaid on the high-resolution X-ray crystal structure of DES-4884 (gray). The oxygen atom of water is represented by a red sphere (simulation) or by a grey sphere (crystal structure; partially obscured by the red sphere). (e) The binding mode of DES-6016, a variant of DES-4884, resolved by high-resolution X-ray crystallography. In all structures, green dashed lines are used to highlight hydrogen bonds.

The crystal structure that includes DES-4799 overlaps with the dominant bound pose in simulation (the heavy-atom RMSD of the pocket is 0.99 Å, and that of the fragment is 0.57 Å). The fragment pose observed in the crystal structure suggested two plausible directions in which the fragment could be extended, so we synthesized ethyl variants (DES-5742 and DES-5743) of the original DES-4799 fragment and obtained high-resolution X-ray structures. These structures reproduce the binding mode of the original aniline fragment.

The crystal structure that includes DES-4884 overlaps well with the structure of the dominant bound pose in MD simulation (the heavy-atom RMSD of the pocket residues is 0.49 Å, and the heavy-atom RMSD of the fragment is 0.70 Å). The crystal structure recovers the observed hydrogen bond to Asn193 and the reorientation of the Phe196 and Phe280 side chains, as seen in the MD simulations (Movie S3). We synthesized DES-6016, a hydroxymethyl derivative of the DES-4884 fragment that extends beyond the hydrogen bond to Asn193 in the direction covered by the Sunesis compound.^22^ The high-resolution crystal structure of DES-6016 bound to PTP1b has the core of the fragment in essentially the same pose as in the crystal structure of DES-4884, adding further experimental support to the DES-4884 binding mode predicted by simulation.

In this communication, we have demonstrated that MD simulation can discover binding poses of fragments at allosteric sites, which were previously unknown, of a challenging drug discovery target. Our simulations incorporated no *a priori* information about possible binding sites. The simulations were also able to sample possible conformational rearrangements of the fragments and the protein during the binding process. The two fragments bound at very different locations, and high-resolution crystal structures of each of these fragments and their variants strongly support the binding modes observed in simulation.

Significantly, one of the binding pockets was occluded by two aromatic side chains in the crystal structure used to initiate the MD simulations, and the open pocket that resulted from the rearrangement of these side chains is rarely observed in published crystal structures. Our results also show that MD simulation can be used to predict binding sites and poses even when fragments have very weak binding strength: The aniline fragment was computationally predicted to have a very weak affinity (>3 mM), and yet the dominant binding pose in our simulations matched well the pose in the subsequent crystal structure. Perhaps counterintuitively, long-timescale MD can be particularly useful when simulating weakly binding fragments, as the long simulations enable a sampling of many different binding modes and multiple reversible binding events at the dominant site.

MD simulations can serve as a tool for obtaining atomically detailed insights about binding modes of weak allosteric binders—a task that is difficult to achieve using traditional biophysical methods. One remaining uncertainty lies in the accuracy with which the underlying physical models used in such simulations (force fields) describe the protein and the small-molecule fragments. With the progress that has been made in the development of force fields, this source of uncertainty has now become manageable for proteins with well-resolved folded-state structures and for fragment-sized small molecules. We anticipate that with the availability of increased computational power and further improvements in force field accuracy, the use of long-timescale, unbiased MD simulation will become increasingly common in fragment-hit validation.

