## Supplementary Information for "Discovery and validation of the binding poses of allosteric fragment hits to PTP1b: From molecular dynamics simulations to X-ray crystallography"

### Supplementary Figures

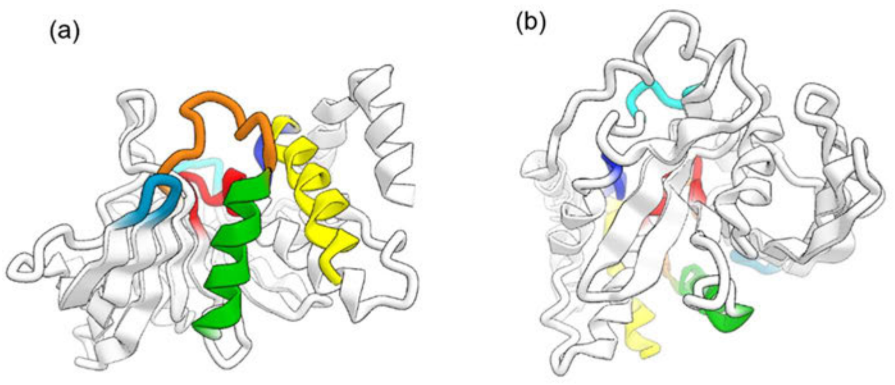


(c)

| DOMAINS | RESIDUES | SEQUENCE |
|---|---|---|
| LOOP | | |
| WPD | T177 to P188 | TTWPDFGVPESP |
| P (or PTP) | H214 to G223 | HCSAGIGRSG |
| Q | R257 to T263 | RMGLIQT |
| L11 | K150 to Y153 | KSYY |
| SBL | N44 to V49 | NRYRDV |
| E | M109 to M133 | MLNRVMEKGSLKCAQ YWPQKEEKEM |
| HELIX | | |
| α3 | A189 to S201 | ASFLNFLFKVRES |
| α6 | D265 to I281 | DQLRFSYLAVIEGAKFI |

**Figure S1**. **Nomenclature of PTP1b regions**. (a) and (b) show two different views of PTP1b, with notable regions colored according to the scheme in (c).

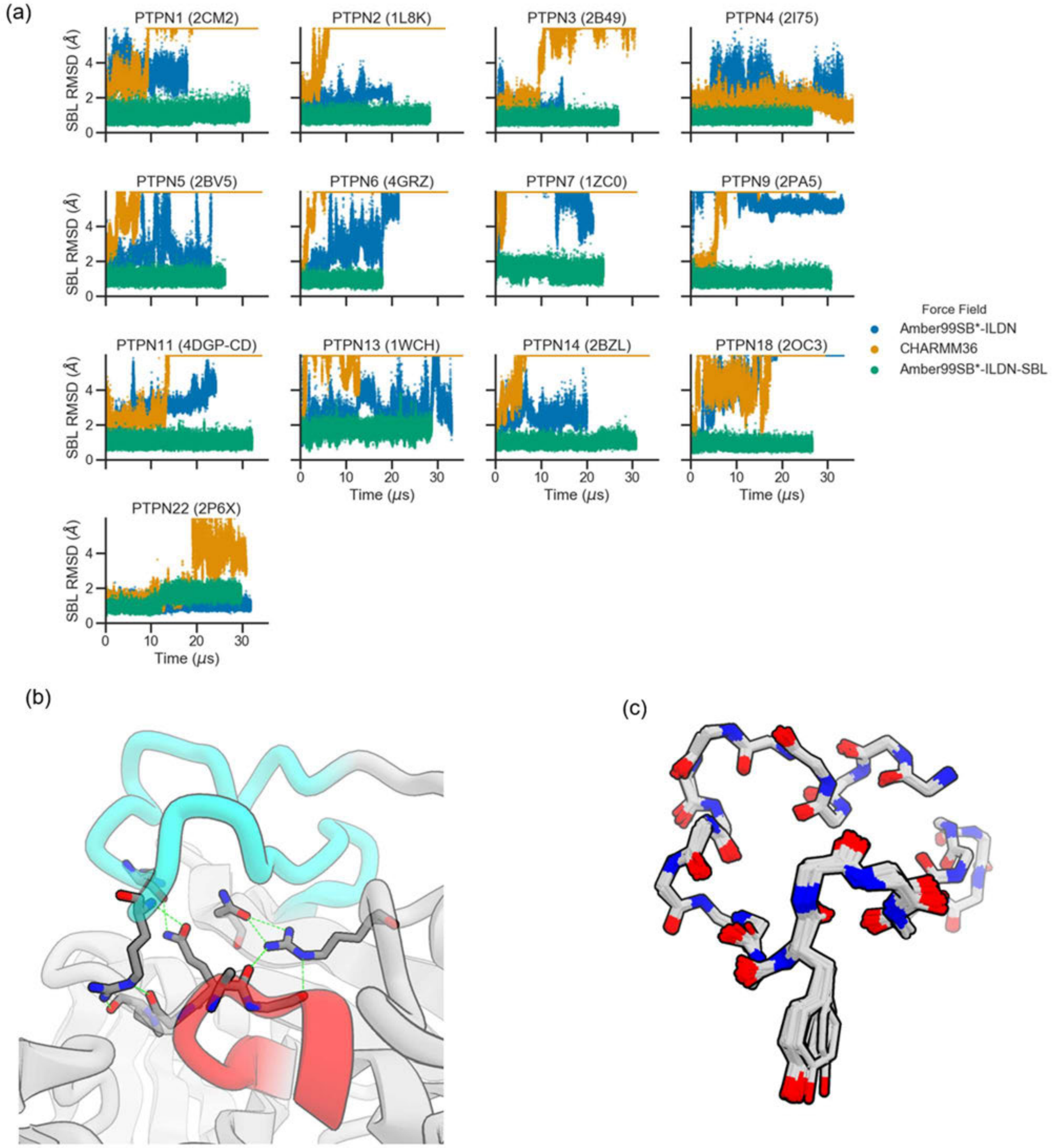


**Figure S2**. The substrate-binding loop (SBL) of non-receptor PTPs is not stable at microsecond timescales in common force fields. (a) MD simulations of non-receptor PTPs were run in

Amber99SB*-ILDN, CHARMM36, and Amber99SB*-ILDN with restraints to stabilize the SBL. Cα-RMSD of the SBL after alignment to a conserved selection from the core of the PTP fold is high (>5 Å) for simulations with Amber99SB*-ILDN and CHARMM36, but low for simulations with SBL restraints. (b) Hydrogen bonds included in the SBL restraints shown with green dashed lines. (c) Overlay of the backbone atoms of the SBL for all deposited PTP1b structures with wild-type sequence.

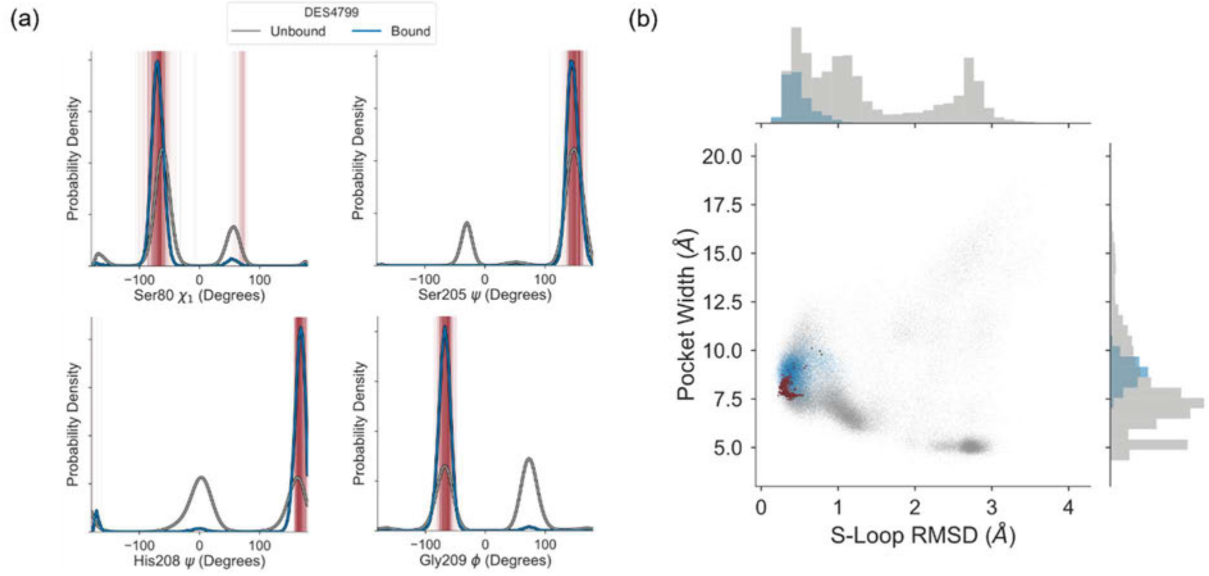


**Figure S3**. Dynamics of the binding pocket stabilized upon DES4799 binding. (a) DES4799 binding stabilizes the primary conformations of key hydrogen bond partner residues observed in available crystal structures (red lines). (b) DES4799 binding stabilizes the conformation of the S-Loop (residues 201–210) to that observed in available crystal structures (red circles). The pocket width is defined as the distance between the $C_\alpha$ atoms of Arg79 and Ser205, and the S-Loop RMSD is computed relative to chain A of the DES4799 co-crystal structure using the $C_\alpha$ atoms of residues 201–210.

### Experimental procedures

All commercially available reagents and solvents were used as received. Reactions using air- or moisture-sensitive reagents were performed under an atmosphere of nitrogen using freshly opened solvents. Reaction progress was monitored by TLC on pre-coated TLC glass plates (silica gel 10–40 µm F254, 1 mm thickness) or by LC/MS (50 mm × 3 mm, 2 µm column; 0.3 µL injection; 5% to 95% MeCN + 0.05% TFA/water + 0.05% TFA gradient over 2 min; 1.2 mL $min^{-1}$ flow; ESI; positive ion mode; UV detection at 220 nm). Flash column chromatography was performed with Biotage CombiFlash Companion systems using prepacked silica gel columns (40–60 µm particle size Agela Technologies columns or similar columns from other vendors). Preparative reverse phase HPLC purifications were typically performed using a XBridge C18 OBD (19 mm × 250 mm; 10 µm stationary phase, with 20 mmol $L^{-1}$ aqueous $NH_4HCO_3$ in acetonitrile gradients as the mobile phase (typically 35–85% acetonitrile over 9 min) with a flow rate of 20 mL $min^{-1}$. NMR spectra were measured on Bruker 300 or 400 MHz spectrometer, and chemical shifts were reported in ppm downfield from TMS using residual nondeuterated solvent as internal standards ($CHCl_3$, 7.26 ppm; DMSO, 2.50 ppm; MeOH, 3.31 ppm). The following abbreviations are used: br = broad signal, s = singlet, d = doublet, dd = doublet of doublets, t = triplet, q = quartet, m = multiplet. The purity of final compounds was verified by LC–MS (3 min run) and $^1H$ NMR (Bruker 300, 400, or 500 MHz spectrometer) to be ≥95% in all cases.

***Synthesis of 3,5-dimethyl-1-(4-nitrophenyl)-1H-pyrazole (DES-4799)***

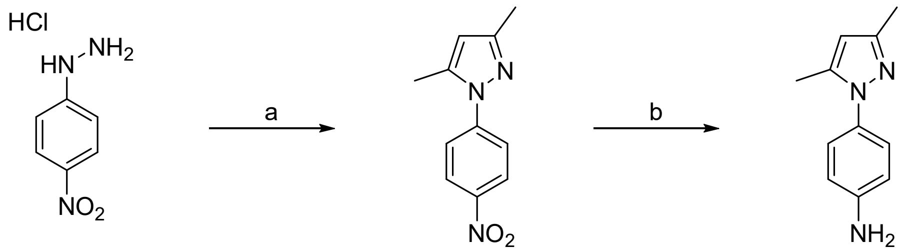

**DES-4799**

Step A: (4-nitrophenyl)hydrazine hydrochloride (1 g, 5 mmol) was dissolved in 6 mL of $H_2O$/Acetic acid (1/1), followed by addition of acetylacetone (0.52 g, 5 mmol). The solution was refluxed for 1 h. The reaction was monitored by LC-MS. pH of solution was basified to 10 by adding 10% NaOH solution. The mixture was diluted with 40 mL ethyl acetate, 10 mL water. Two phases were separated, the organic layer was washed with water and brine, dried ($Na_2SO_4$), and concentrated. The crude was purified by normal phase silica column (H:EA 100:0 to 75:25) to afford 3,5-dimethyl-1-(4-nitrophenyl)-1H-pyrazole (0.13 g, 12%). LCMS (ESI) calculated for $C_{11}H_{11}N_3O_2$ $[M + H]^+$: 218.1, found 218.1; $^1$H-NMR (500 MHz, $CDCl_3$) δ 8.30 (d, $J$ = 9.2 Hz, 2H), 7.66 (d, $J$ = 9.2 Hz, 2H), 6.07 (s, 1H), 2.42 (s, 3H), 2.29 (s, 3H).

Step B: 3,5-dimethyl-1-(4-nitrophenyl)-1H-pyrazole (40 mg, 0.18 mmol) was dissolved in 1 mL ethyl acetate, and 3 mL MeOH. 50 mg Pd/C (10%, wet) was added, the flask was purged with hydrogen and the reaction was carried out under hydrogen balloon pressure overnight at room temperature. The catalyst was removed using celite pad, and the solvent was evaporated to get 4-(3,5-dimethyl-1H-pyrazol-1-yl)aniline (33 mg, 98%). LCMS (ESI) calculated for $C_{11}H_{13}N_3$ $[M + H]^+$: 188.1, found 188.2; $^1$H-NMR (500 MHz, $CDCl_3$) δ 7.14 (d, $J$ = 8.6 Hz, 2H), 6.68 (d, $J$ = 8.6 Hz, 2H), 5.93 (s, 1H), 3.78 (s, br, 2H), 2.27 (s, 3H), 2.20 (s, 3H).

***Synthesis of 4-(3-ethyl-5-methyl-1H-pyrazol-1-yl)aniline (DES-5742) and 4-(5-ethyl-3-methyl-1H-pyrazol-1-yl)aniline (DES-5743)***

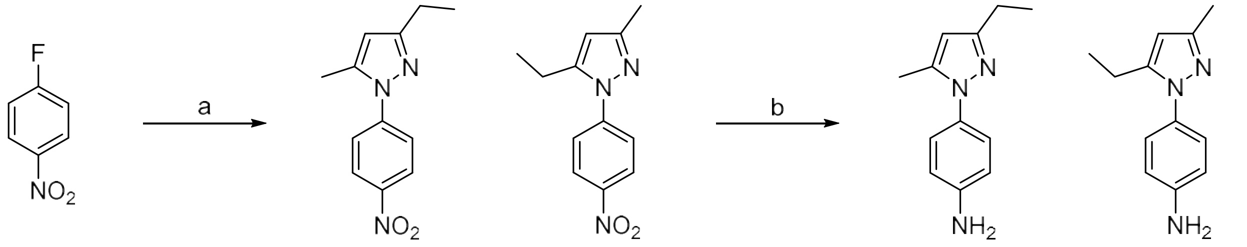

1 2 DES-5742 DES-5743

Step A: 1-fluoro-4-nitrobenzene (1.27 g, 9 mmol), 3-ethyl-5-methyl-1H-pyrazole (1 g, 9 mmol) and KOtBu (1.1 g, 9.9 mmol) were mixed in a MW tube (solvent-free) and heated at 70 ºC for 8 min. Ethyl acetate (40 mL) was added and the solution was transferred to a separating funnel. The organic layer was washed with water, dried ($Na_2SO_4$), and concentrated. The crude was purified flash chromatography, normal phase (H:EA 100:0 to 100:15), to obtain two regioisomers (181 mg compound 1, and 30 mg compound 2).

3-ethyl-5-methyl-1-(4-nitrophenyl)-1H-pyrazole (Compound 1): LCMS (ESI) calculated for $C_{12}H_{13}N_3O_2$ $[M + H]^+$: 232.1, found 232.3, $^1$H-NMR (500 MHz, $CDCl_3$) δ 8.31 (d, $J$ = 9.1 Hz, 2H), 7.68 (d, $J$ = 9.1 Hz, 2H), 6.10 (s, 1H), 2.67 (q, $J$ = 7.6 Hz, 2H), 2.43 (s, 3H), 1.27 (t, $J$ = 7.6 Hz, 3H).

5-ethyl-3-methyl-1-(4-nitrophenyl)-1H-pyrazole (Compound 2): LCMS (ESI) calculated for $C_{12}H_{13}N_3O_2$ $[M + H]^+$: 232.1, found 232.3, $^1$H-NMR (500 MHz, $CDCl_3$) δ 8.31 (d, $J$ = 9.1 Hz, 2H), 7.65 (d, $J$ = 9.1 Hz, 2H), 6.11 (s, 1H), 2.75 (q, $J$ = 7.5 Hz, 2H), 2.31 (s, 3H), 1.27 (t, $J$ = 7.5 Hz, 3H).

Step B: 3-Ethyl-5-methyl-1-(4-nitrophenyl)-1H-pyrazole (100 mg, 0.43 mmol) was dissolved in 0.5 mL EA, and 5 mL MeOH. 100 mg Pd/C (10%, wet) was added, the flask was purged with hydrogen, and reaction was carried out under hydrogen balloon pressure overnight at room temperature. Catalyst was removed using celite pad, and solvent was evaporated. The crude was purified by flash chromatography, normal phase (H:EA 100:0 to 100:50), to obtain 4-(3-ethyl-5-methyl-1H-pyrazol-1-yl)aniline (24 mg, 28%). LCMS (ESI) calculated for $C_{12}H_{15}N_3$ $[M + H]^+$: 202.1, found 202.2, $^1H$ NMR (500 MHz, DMSO-$d_6$) δ 7.04 (d, $J$ = 8.6 Hz, 2H), 6.61 (d, $J$ = 8.7 Hz, 2H), 5.98 (s, 1H), 5.27 (s, 2H), 2.56 – 2.45 (m, 2H), 2.16 (s, 3H), 1.16 (t, $J$ = 7.6 Hz, 3H).

Step B: 5-ethyl-3-methyl-1-(4-nitrophenyl)-1H-pyrazole (30 mg, 0.13 mmol) was dissolved in 0.5 mL EA, and 5 mL MeOH. 50 mg Pd/C (10%, wet) was added, the flask was purged with hydrogen and reaction was carried out under hydrogen balloon pressure overnight at room temperature. Catalyst was removed using celite pad, and solvent was evaporated. The crude was purified by flash chromatography, normal phase (H:EA 100:0 to 100:50), to obtain 4-(5-ethyl-3-methyl-1H-pyrazol-1-yl)aniline (12 mg, 47%) LCMS (ESI) calculated for $C_{12}H_{15}N_3$ $[M + H]^+$: 202.1, found 202.2, $^1H$-NMR (500 MHz, DMSO-$d_6$) δ 7.00 (d, $J$ = 8.6 Hz, 2H), 6.60 (d, $J$ = 8.7 Hz, 2H), 5.96 (s, 1H), 5.29 (s, 2H), 2.53 – 2.45 (m, 2H), 2.14 (s, 3H), 1.08 (t, $J$ = 7.5 Hz, 3H).

***Synthesis of 6-methyl-4-(piperazin-1-yl)-2-(trifluoromethyl)quinoline (DES-4884)***

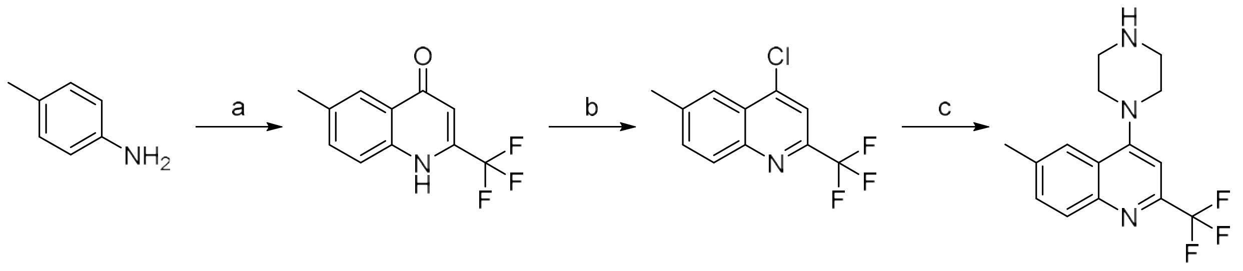

Step A: To a mixture of ethyl 4,4,4-trifluoro-3-oxobutanoate (1.72 g, 9.33 mmol) and polyphosphoric acid (7.50 g), was added 4-methylaniline (1.00 g, 9.33 mmol) over 15 min at 100 °C and with mechanical stirring. After the addition, the temperature was then raised to 150 °C and kept constant for 2 h. After cooling to room temperature, the mixture was diluted with aqueous NaOH (5%, 80 mL). The precipitate formed was dissolved in a 10% aqueous NaOH (40 mL). After some insoluble material had been removed by filtration, the clear solution was acidified with 30% HCl. The solid was collected and crystallized from ethanol. The crystal was purified by silica gel column chromatography, eluted with (EA/PE = 1/5) to afford 6-methyl-2-(trifluoromethyl)quinolin-4(1*H*)-one (0.40 g, 19%) as a light yellow solid: LCMS (ESI) calculated for $C_{11}H_8F_3NO$ $[M + H]^+$: 228, found 228; $^1$H-NMR (400 MHz, DMSO-$d_6$) δ 12.13 (br, 1H), 7.96 (s, 1H), 7.88 (br, 1H), 7.61 (d, $J$ = 8.5 Hz, 1H), 7.12 (br, 1H), 2.48 (s, 3H); $^{19}$F-NMR (376 MHz, DMSO-$d_6$) δ -66.48.

Step B: To a solution of 6-methyl-2-(trifluoromethyl)-1,4-dihydroquinolin-4-one (0.10 g, 0.44 mmol) and N,N-dimethylaniline (0.10 g, 0.88 mmol) in toluene (5 mL) was added POCl3 (0.13 g, 0.88 mmol) at room temperature. The reaction solution was allowed to warm to 90 °C and stirred for 4 h. After cooling to room temperature, the reaction was quenched with water (20 mL), and then extracted with DCM (3 x 20 mL). The combined organic layers were washed with brine (2 x 15 mL), dried over anhydrous $Na_2SO_4$ and filtered. The filtrate was concentrated under vacuum. The residue was purified by silica gel column chromatography, eluted with EA/PE (1/3) to afford 4-chloro-6-methyl-2-(trifluoromethyl)quinoline (90 mg, 88%) as a greenish oil: LCMS (ESI) calculated for $C_{11}H_7ClF_3N$ $[M + H]^+$: 246, 248 (3 : 1), found 246, 248 (3 : 1); 1H-NMR (300 MHz, DMSO-$d_6$) δ 8.19 (s, 1H), 8.12 (d, $J$ = 8.6 Hz, 1H), 8.05 (s, 1H), 7.85 (d, $J$ = 8.6 Hz, 1H), 2.60 (s, 3H).

Step C: 4-chloro-6-methyl-2-(trifluoromethyl)quinoline (100 mg, 0.41 mmol) was dissolved in DMF (3 mL). Piperazine (71 mg, 0.82 mmol) and $K_2CO_3$ (166 mg, 1.2 mmol) were added and the reaction mixture was stirred at 120°C for 16 h. The reaction mixture was then transferred to a separating funnel, diluted with ethyl acetate, washed with brine, and finally dried with anhydrous $Na_2SO_4$. The solvent was removed under vacuum to obtain 6-methyl-4-(piperazin-1-yl)-2-(trifluoromethyl)quinoline (62 mg, 51 %). LCMS (ESI) calculated for $C_{12}H_{15}N_3$ $[M + H]^+$: 296.1, found 296.2, $^1$H-NMR (500 MHz, DMSO) δ 7.96 (d, $J$ = 8.6 Hz, 1H), 7.83 (s, 1H), 7.66 (dd, $J$ = 8.6, 1.8 Hz, 1H), 7.19 (s, 1H), 3.23 – 3.14 (m, 4H), 3.02 – 2.95 (m, 4H), 2.55 (s, 3H).

***Synthesis of (4-(6-methyl-2-(trifluoromethyl)quinolin-4-yl)piperazin-2-yl)methanol (DES-6016)***

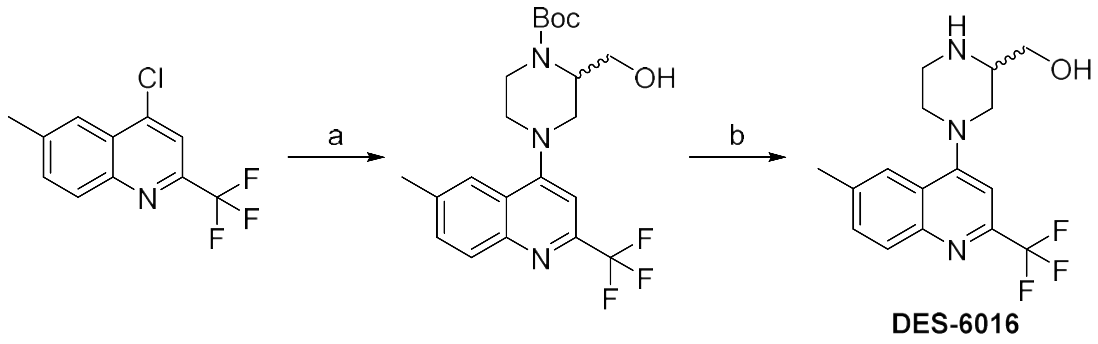

Step A: To a solution of *tert*-butyl 2-(hydroxymethyl)piperazine-1-carboxylate (0.85 g, 3.93 mmol) and DIEA (0.80 g, 6.19 mmol) in NMP (5 mL) was added 4-chloro-6-methyl-2-(trifluoromethyl)quinolone (0.50 g, 2.04 mmol) at room temperature. The reaction solution was allowed to warm to 120 °C and stirred for 4 h. The reaction was diluted with water (30 mL) and extracted with EA (3 x 50 mL). The combined organic layers were washed with brine (2 x 15 mL), dried over anhydrous $Na_2SO_4$, and filtered. The filtrate was concentrated under vacuum. The residue was purified by silica gel column chromatography, eluted with PE/EA (3/2) to afford *tert*-butyl 2-(hydroxymethyl)-4-[6-methyl-2-(trifluoromethyl)quinolin-4-yl]piperazine-1-carboxylate (0.35 g, 40%) as a light yellow semi-solid: LCMS (ESI) calculated for $C_{21}H_{26}F_3N_3O_3$ $[M + H]^+$: 426, found 426; $^1$H-NMR (400 MHz, $CDCl_3$) δ 8.13 (d, *J* = 8.6 Hz, 1H), 7.91 (s, 1H), 7.62 (d, *J* = 8.6, 1H), 7.16 (s, 1H), 4.42 (s, 1H), 4.21-4.05 (m, 3H), 3.86-3.82 (m, 1H), 3.60-3.44 (m, 3H), 3.12-2.94 (m, 2H), 2.60 (s, 3H), 1.54 (s, 9H).

Step B: A solution of *tert*-butyl 2-(hydroxymethyl)-4-[6-methyl-2-(trifluoromethyl)quinolin-4-yl]piperazine-1-carboxylate (50 mg, 0.15 mmol) in TFA (0.5 mL) and DCM (1 mL) was stirred at room temperature for 2 h. The reaction solution was concentrated under vacuum. The residue was purified by Prep-HPLC with the following conditions: Column: XBridge C18 OBD, 100 Å, 10 μm, 19 mm x 250 mm; Mobile Phase A: water (20 mmol $L^{-1}$ $NH_4HCO_3$), Mobile Phase B: ACN; Flow rate: 20 mL $min^{-1}$; Gradient: 35% B to 85% B in 9 min; Detector: UV 254/210 nm; Retention time: 8.54 min. The collected fractions containing the desired product were concentrated and lyophilized to afford (4-(6-methyl-2-(trifluoromethyl)quinolin-4-yl) piperazin-2-yl)methanol (15 mg, 39%) as an off-white solid: LCMS (ESI) calculated for $C_{16}H_{18}F_3N_3O$ $[M + H]^+$: 326, found 326; $^1$H-NMR (400 MHz, $CD_3OD$) δ 7.99 (d, *J* = 8.4 Hz, 1H), 7.91 (s, 1H), 7.66 (dd, *J* = 8.4, 1.6 Hz, 1H), 7.23 (s, 1H), 3.68-3.56 (m, 4H), 3.23-3.17 (m, 3H), 3.04-2.97 (m, 1H), 2.80-2.75 (m, 1H), 2.58 (s, 3H); $^{19}$F-NMR (376 MHz, $CD_3OD$) δ -69.05.

### SPR Assay

The SPR assay was outsourced to Biosensor Tools, LLC.[1] PTP1b (residues 5–321) and five off-target proteins (CAII, SA, Ovalbumin, IgG, and superoxide dismutase) were immobilized on CM7 sensor chips in Biacore. The instrument temperature was dropped to 4 ºC. The compounds' binding was tested to each of the six protein surfaces at 300 μM concentration. Running buffer included 10 nM HEPES, 150 nM NaCl, 0.005% p20, 0.5 mM TCEP, and 1% DMSO. Two plates of the Maybridge fragment library were tested at 100 μL $min^{-1}$ with 11-second association and dissociation time.

DES-4884 was the most selective molecule from the set of molecules included in Maybridge plate 1. This compound still showed some background binding to the control protein surfaces. A $K_d$ estimate for DES-4884 binding to PTP1b was ~2 mM at 4 ºC.

DES-4799 was the most selective for PTP1b of all the compounds in plate 2 of the Maybridge library. This compound still showed some background binding to the control protein surfaces. A $K_d$ estimate for DES-4799 to PTP1b was 1.2 mM at 4 ºC.

### Crystallography

The crystallization conditions, data collection statistics, and refined structures have been deposited to the PDB. The PTP1b complexes with the respective fragments have the following PDB IDs: 8G65 (DES4799), 8G67 (DES4884), 8G68 (DES5742), 8G69 (DES5743), and 8G6A (DES6016).

### Simulation Details

All simulations were run on Anton,[2] a specialized machine for molecular dynamics simulations.

In order to test the stability of the substrate binding loop (SBL) in classical PTPs, protein models for PTPN1 (PDB: 2CM2), PTPN2 (PDB: 1L8K), PTPN3 (PDB: 2B49), PTPN4 (PDB: 2I75), PTPN5 (PDB: 2BV5), PTPN6, (PDB: 4GRZ), PTPN7 (PDB: 1ZC0), PTPN9 (PDB: 2PA5), PTPN11 (PDB: 4DGP), PTPN13 (PDB: 1WCH), PTPN14 (PDB: 2BZL), PTPN18 (PDB: 2OC3), and PTPN22 (PDB: 2P6X) were prepared using SWISS-MODEL to fill in missing loops. For PTPN1, the N-terminal His tag was removed and the three missing Gly283 and Asp284 were added to result in a 2CM2(2-284) construct. For PTPN11, the full-length construct including the N-SH2 and C-SH2 was prepared, and then truncated to the PTP domain (residues 217–528).

We tested Amber ff99SB*-ILDN[3–5] and CHARMM36[6] protein force fields on all constructs and discovered that it is necessary to impose additional restraints on the SBL to maintain structural integrity of each protein construct (Figure S2A). The SBL restraints used to enhance the stability of the SBL in simulation were determined based on conserved motifs across the classical PTPs, in order to ensure transferability across PTPs. The restraints involved the application of a harmonic potential of 1 kcal $mol^{-1}$ $radian^{-1}$ to the backbone dihedrals of the SBL residues (Figure S2B, S2C), and a set of distance restraints (Table S1) to enforce the hydrogen bond network that is conserved among all classical PTPs. The backbone dihedral restraints were applied around the Phi/Psi values present in the crystal structure for the PTPs to the SBL residues in order to stabilize the observed conformations (Table S2).

The protein constructs in simulations of fragments with PTPN1 using DES4799 and DES4884 were prepared similarly to PTPN1 apo simulations. The proteins were parameterized by Amber ff99SB*-ILDN with the SBL restraints. The ligands were placed randomly and parameterized by the generalized Amber force field (GAFF).[7]

Protein and protein-ligand systems were then solvated in a 90 × 90 × 90 $Å^3$ cubic box of 150 mM NaCl solution, each containing ~63,000 atoms. Each system was equilibrated in the NPT ensemble for 50 ns with harmonic position restraints on all heavy protein atoms, tapered linearly to 0 from 5 kcal $mol^{-1}$ $Å^{-2}$ . Production runs were subsequently performed in the NVT ensemble from the final frame of the NPT relaxation simulation by coupling the system to a Nosé-Hoover thermostat[8,9] at 310 K with a relaxation time of 1 ps. A RESPA integrator[10] was used with a time step of 2.4 fs. The long-range electrostatic forces were calculated in k-space using a grid-based method with Gaussian spreading to the grid every 7.5 fs.[11]

### Tables

| PTPN | ATOM1 | ATOM2 |
|---|---|---|
| 1 | ARG45 NE | GLY86 O |
| 1 | ARG45 NH2 | PRO87 O |
| 1 | GLN85 NE2 | ARG43 O |
| 1 | GLN85 OE1 | ARG45 N |
| 1 | ARG257 NH2 | ALA217 O |
| 1 | ARG257 NE | GLY218 O |
| 1 | ARG257 NH2 | ASN68 OD1 |
| 1 | ARG257 NH1 | ASN68 OD1 |
| 2 | ARG47 NE | GLY88 O |
| 2 | ARG47 NH2 | PRO 89 O |
| 2 | GLN87 NE2 | ARG47 O |
| 2 | GLN87 OE1 | ARG47 N |
| 2 | ARG255 NH2 | ALA218 O |
| 2 | ARG255 NE | GLY219 O |
| 2 | ARG255 NH2 | ASN70 OD1 |

| | | |
|---|---|---|
| 2 | ARG255 NH1 | ASN70 OD1 |
| 3 | ARG675 NE | GLY719 O |
| 3 | ARG675 NH2 | PRO 720 O |
| 3 | GLN718 NE2 | LYS673 O |
| 3 | GLN718 OE1 | ARG675 N |
| 3 | ARG881 NH2 | ALA844 O |
| 3 | ARG881 NE | GLY845 O |
| 3 | ARG881 NH2 | ASN697 OD1 |
| 3 | ARG881 NH1 | ASN697 OD1 |
| 4 | ARG684 NE | GLY728 O |
| 4 | ARG684 NH2 | PRO729 O |
| 4 | GLN727 NE2 | LYS682 O |
| 4 | GLN727 OE1 | ARG684 N |
| 4 | ARG891 NH2 | ALA854 O |
| 4 | ARG891 NE | GLY855 O |
| 4 | ARG891 NH2 | ASN706 OD1 |

| | | |
|---|---|---|
| 4 | ARG891 NH1 | ASN706 OD1 |
| 5 | ARG303 NE | GLY350 O |
| 5 | ARG303 NH2 | PRO351 O |
| 5 | GLN349 NE2 | LYS301 O |
| 5 | GLN349 OE1 | ARG303 N |
| 5 | ARG511 NH2 | ALA473 O |
| 5 | ARG511 NE | GLY474 O |
| 5 | ARG511 NH2 | ASN331 OD1 |
| 5 | ARG511 NH1 | ASN331 OD1 |
| 6 | ARG275 NE | GLY326 O |
| 6 | ARG275 NH2 | CYS327 O |
| 6 | GLN325 NE2 | LYS273 O |
| 6 | GLN325 OE1 | ARG275 N |
| 6 | ARG495 NH2 | ALA455 O |
| 6 | ARG495 NE | GLY456 O |
| 6 | ARG495 NH2 | ASN303 OD1 |

| | | |
|---|---|---|
| 6 | ARG495 NH1 | ASN303 OD1 |
| 7 | ARG103 NE | GLY149 O |
| 7 | ARG103 NH2 | PRO150 O |
| 7 | GLN148 NE2 | LYS101 O |
| 7 | GLN148 OE1 | ARG103 N |
| 7 | ARG309 NH2 | ALA272 O |
| 7 | ARG309 NE | GLY273 O |
| 7 | ARG309 NH2 | ASN130 OD1 |
| 7 | ARG309 NH1 | ASN130 OD1 |
| 9 | ARG332 NE | GLY377 O |
| 9 | ARG332 NH2 | PRO378 O |
| 9 | GLN376 NE2 | LYS330 O |
| 9 | GLN376 OE1 | ARG332 N |
| 9 | ARG554 NH2 | ALA517 O |
| 9 | ARG554 NE | GLY518 O |
| 9 | ARG554 NH2 | ASN359 OD1 |

| | | |
|---|---|---|
| 9 | ARG554 NH1 | ASN359 OD1 |
| 11 | ARG278 NE | GLY332 O |
| 11 | ARG278 NH2 | CYS333 O |
| 11 | GLN331 NE2 | LYS276 O |
| 11 | GLN331 OE1 | ARG278 N |
| 11 | ARG501 NH2 | ALA461 O |
| 11 | ARG501 NE | GLY462 O |
| 11 | ARG501 NH2 | ASN306 OD1 |
| 11 | ARG501 NH1 | ASN406 OD1 |
| 13 | ARG2242 NE | GLY2284 O |
| 13 | ARG2242 NH2 | PRO2285 O |
| 13 | GLN2283 NE2 | LYS2240 O |
| 13 | GLN2284 OE1 | ARG2242 N |
| 13 | ARG2447 NH2 | ALA2410 O |
| 13 | ARG2447 NE | GLY2411 O |
| 13 | ARG2447 NH2 | ASN2264 OD1 |

| | | |
|---|---|---|
| 13 | ARG2447 NH1 | ASN2264 OD1 |
| 14 | ARG938 NE | GLY984 O |
| 14 | ARG938 NH2 | PRO985 O |
| 14 | GLN983 NE2 | ARG936 O |
| 14 | GLN983 OE1 | ARG938 N |
| 14 | ARG1160 NH2 | ALA1123 O |
| 14 | ARG1160 NE | GLY1124 O |
| 14 | ARG1160 NH2 | ASN964 OD1 |
| 14 | ARG1160 NH1 | ASN964 OD1 |
| 18 | ARG61 NE | GLY106 O |
| 18 | ARG61 NH2 | PRO107 O |
| 18 | GLN105 NE2 | LYS59 O |
| 18 | GLN105 OE1 | ARG61 N |
| 18 | ARG271 NH2 | ALA231 O |
| 18 | ARG271 NE | GLY232 O |
| 18 | ARG271 NH2 | ASN88 OD1 |

| | | |
|---|---|---|
| 18 | ARG271 NH1 | ASN88 OD1 |
| 22 | ARG59 NE | GLY104 O |
| 22 | ARG59 NH2 | PRO105 O |
| 22 | GLN103 NE2 | LYS57 O |
| 22 | GLN103 OE1 | ARG59 N |
| 22 | ARG269 NH2 | ALA229 O |
| 22 | ARG269 NE | GLY230 O |
| 22 | ARG269 NH2 | ASN86 OD1 |
| 22 | ARG269 NH1 | ASN86 OD1 |

**Table 1.** Distance restraints of the SBL that enforce the hydrogen bond network. The first column shows the gene, and the second and third columns show the pair of atoms that form the hydrogen bond.

| PTPN | RESIDUES |
|---|---|
| 1 | 32-56 |
| 2 | 34-58 |
| 3 | 662-686 |
| 4 | 671-695 |
| 5 | 314-338 |
| 6 | 262-286 |
| 7 | 111-135 |
| 8 | 319-343 |
| 11 | 265-289 |
| 13 | 2229-2253 |
| 14 | 925-949 |
| 18 | 48-72 |
| 22 | 46-70 |

**Table 2.** Backbone dihedral restraints of the SBL. The first column shows the gene, and the second column the range of residues that is restrained.
